# Phase separation of Ede1 promotes the initiation of endocytic events

**DOI:** 10.1101/861203

**Authors:** Mateusz Kozak, Marko Kaksonen

## Abstract

Clathrin-mediated endocytosis is a major pathway that eukaryotic cells use to produce transport vesicles from the plasma membrane. The assembly of the endocytic coat is initiated by a dynamic network of weakly interacting proteins, but the exact mechanism of initiation is unknown. Ede1, the yeast homologue of mammalian Eps15, is one of the early-arriving endocytic proteins and a key initiation factor. In the absence of Ede1, most other early endocytic proteins lose their punctate localization and the frequency of endocytic initiation is decreased. We show here that in mutants with increased amounts of cytoplasmic Ede1, the excess protein forms large condensates which exhibit properties of phase separated liquid protein droplets. These Ede1 condensates recruit many other early-arriving endocytic proteins. Their formation depends on the core region of Ede1 that contains a coiled coil and a low-complexity domain. We demonstrate that Ede1 core region is essential for the endocytic function of Ede1. The core region can also promote clustering of a heterologous lipid-binding domain into discrete sites on the plasma membrane that initiate endocytic events. We propose that the clustering of the early endocytic proteins and cargo depend on phase separation mediated by Ede1.

## Introduction

Clathrin-mediated endocytosis is the primary route of specific internalization of extracellular and surface molecules in eukaryotic cells (Kirchhausen et al., 2014). This process depends on a complex protein machinery that assembles in a sequential manner on the plasma membrane (Kaksonen et al., 2005; Sirotkin et al., 2010; Taylor et al., 2011). An endocytic event starts with the assembly of a group of conserved pioneer proteins that select the site and initiate the assembly of endocytic proteins (Kaksonen and Roux, 2018). These proteins include membrane binding adaptor proteins, such as the AP2-complex, Syp1 (FCHo1/2 in mammals), and Yap1801/2 (AP180), and scaffold proteins, such as clathrin and Ede1 (Eps15). Different pioneer proteins (Cocucci et al., 2012; Henne et al., 2010; Ma et al., 2016), lipids (Antonescu et al., 2011) and cargo molecules (Layton et al., 2011; Liu et al., 2010) have been shown to promote the initiation step (reviewed by Godlee and Kaksonen, 2013). However, the exact mechanism by which the endocytic site assembly is initiated remains poorly understood.

In budding yeast *Saccharomyces cerevisiae*, the initiation of endocytosis is extremely robust and does not depend on any single pioneer protein. As many as seven early endocytic proteins can be simultaneously deleted without completely blocking the endocytic initiation, although the frequency of the endocytic events and the regulation of cargo recruitment is severely compromised (Brach et al., 2014).

One of the key early proteins is Ede1, a homologue of mammalian Eps15. It is among the earliest proteins to appear at the nascent endocytic site (Carroll et al., 2012). A deletion of the *EDE1* gene reduces the overall membrane uptake by 35 % (Gagny et al., 2000) and decreases the number of productive endocytic events by 50 % (Carroll et al., 2012; Kaksonen et al., 2005). Ede1 oligomerizes via its coiled-coil domain, which is required for it to properly localise and function (Boeke et al., 2014; Lu and Drubin, 2017). Ede1 also interacts via its three EH domains and other interaction motifs with several adaptor proteins such as the AP-2 complex, epsins, Yap1801/2, Sla2 and Syp1 (Maldonado-Báez et al., 2008; Reider et al., 2009). These multiple interactions are known to be critical for the ability of Ede1 to locally concentrate other endocytic proteins on the plasma membrane to initiate an endocytic event.

In recent years, phase separation has garnered much attention as a mechanism for assembly of many cellular structures, including signalling clusters of membrane receptors (Case et al., 2019a). Liquid, phase-separated biomolecular condensates also form many cellular organelles which are not bound by membranes, such as P-granules, nucleoi, stress granules and others (Banani et al., 2017; Shin and Brangwynne, 2017). Such membraneless compartments can accelerate reactions, sequester molecules from the cytoplasm or establish spatial organisation.

In this work, we show that Ede1 has a propensity to form cellular condensates that exhibit the characteristics of phase separated liquids. We identify the molecular features driving the condensation of Ede1 and show that they are essential for the normal function of Ede1 in endocytic initiation. Our findings suggest that phase separation of Ede1 may be the mechanisms by which it mediates the initiation of endocytic events.

## Results

### Ede1 can form dynamic protein condensates

In normal yeast cells, fluorescently tagged Ede1 localizes to endocytic sites at the plasma membrane (Kukulski et al., 2012). However, we discovered previously that under certain experimental conditions Ede1 can also assemble into large condensates (Boeke et al., 2014). These condensates were seen in cells that either overexpressed Ede1, or expressed Ede1 at normal levels, but lacked three early endocytic adaptors. Although these condensates are abnormal structures that have not been observed in wild-type cells, we reasoned that studying them in more detail might provide insights into the mechanism by which Ede1 promotes the assembly of the early endocytic proteins.

To visualize the condensates, we expressed Ede1-EGFP from its endogenous locus in haploid wild-type cells, or in cells from which three endocytosis-related genes were deleted (*yap1801Δ yap1802Δ apl3Δ*, called 3×ΔEA for short). These genes code for early arriving endocytic adaptor proteins Yap1801, Yap1802 and the α-subunit of the AP-2 complex. Alternatively, we overexpressed EGFP-Ede1 from its endogenous locus under the control of a strong heterologous promoter.

We observed that part of the cellular Ede1-EGFP in the mutant strains localized into condensates that were much brighter than the normal endocytic sites (Figure 1A). The condensates in the overexpression strain were larger and brighter than those in the 3×ΔEA cells (Figure 1B). The condensates usually associated with the plasma membrane, but were also observed away from it (Figure 1A). In contrast, normal endocytic sites are always associated with the plasma membrane. The condensates in the Ede1 overexpression strain were often large enough that their shape was resolvable (Figure 1B). They appeared circular in surface view, and as dome-like structures limited by the plasma membrane in the side view. The Ede1 condensates were remarkably long lived and we have observed individual condensates for up to one hour (Figure 1–video 1). This stands in contrast with normal endocytic sites, where Ede1-EGFP typically persists for 1 to 2 minutes (Stimpson et al., 2009). Despite their stability, some of the condensates appeared to undergo dynamic fission and fusion events, suggesting that they are not solid aggregates (Figure 1C).

**Figure 1.**
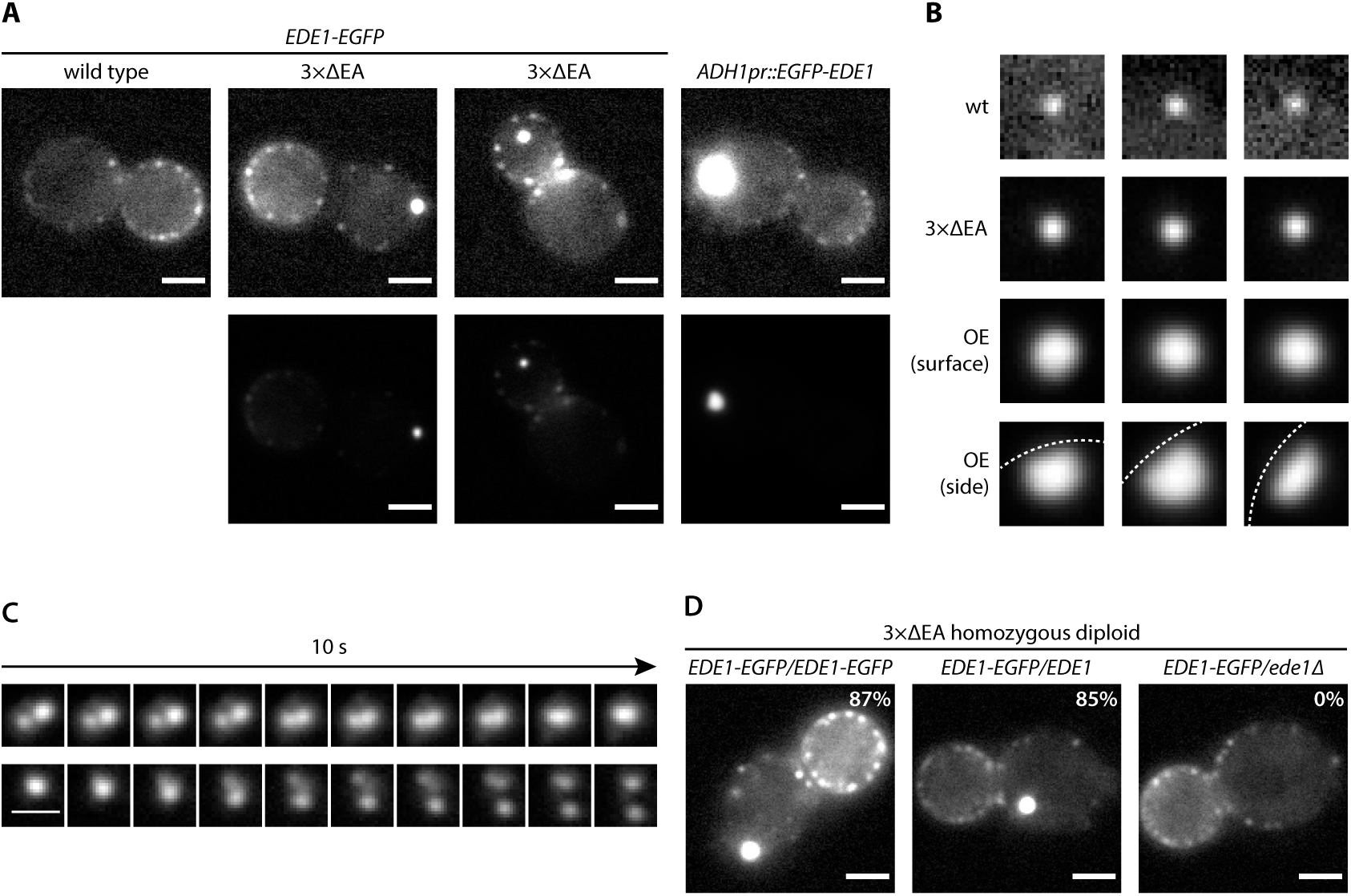
Excess cytosolic Ede1 assembles into condensates in vivo. (A) Representative images of yeast cells expressing Ede1-EGFP in wild-type and 3×ΔEA genetic backgrounds, or overexpressing EGFP-Ede1 under the control of ADH1 promoter. In the top row, mutant cell micrographs are shown using the same display range as the wild type. In the bottom row, the same micrographs are shown using their full display range. Two cells are shown for 3×ΔEA background to display the membrane-associated and cytoplasmic localizations of Ede1 condensates. Scale bars: 2 μm. (B) Representative images of Ede1-EGFP at endocytic sites in wild-type background (wt), Ede1-EGFP condensates in 3×ΔEA cells, and EGFP-Ede1 overexpression-induced condensates (OE). OE condensates are shown in two different orientations. Each frame is 1.5 μm×1.5 μm; dotted white line represents the approximate position of the plasma membrane. (C) Two time series of Ede1-EGFP condensates undergoing apparent fusion (top) and fission (bottom) events. Scale bar: 1 μm. (D) Representative images of Ede1-EGFP in diploid cells homozygous for the 3×ΔEA background and differing in the Ede1 locus: *EDE1-EGFP/EDE1-EGFP, EDE1-EGFP/EDE1* or *EDE1-EGFP/ede1Δ*. Percentages represent the fraction of cells containing condensates in each strain, number of analysed cells between 40 and 47. Scale bars: 2 μm. **Figure 1–video 1.** One-hour movie of Ede1-EGFP in 3×ΔEA cells. Scale bar: 2 μm.

All three adaptors absent from 3×ΔEA cells interact with Ede1, as well as membrane lipids and protein cargo. As Ede1 does not have known membrane-binding activity, it is likely that the cytosolic pool of Ede1 is increased both in the overexpression and the 3×ΔEA backgrounds, and the excess protein assembles into the condensates. To test whether the formation of condensates in 3×ΔEA background depends on Ede1 concentration, we generated diploid cells homozygous for the three adaptor deletions. We expressed Ede1-EGFP in these cells either from both *EDE1* alleles (*EDE1-EGFP/EDE1-EGFP*), or from one allele in combination with untagged *EDE1* (*EDE1-EGFP/EDE1*) or a deletion of *EDE1* (*EDE1-EGFP/ede1Δ*). The condensates formed in both strains expressing two alleles of EDE1, but not in the strain where only one EDE1 allele was present (Figure 1D). This result suggests that the condensate assembly depends on Ede1 concentration.

### Ede1 condensates exhibit liquid-like properties

Because of their spherical shapes, concentration dependence and dynamic behaviours, we hypothesized that the Ede1 condensates might be phase-separated liquid droplets. To test this idea, we first performed fluorescence recovery after photobleaching (FRAP) experiments on Ede1-EGFP condensates in the 3×ΔEA background. After photobleaching, the condensates rapidly recovered most of their fluorescence (Figure 2A). The recovery halftime of a single-exponential FRAP model fitted to an average of 36 events was 22 s, and the mobile fraction was 63 %.

**Figure 2.**
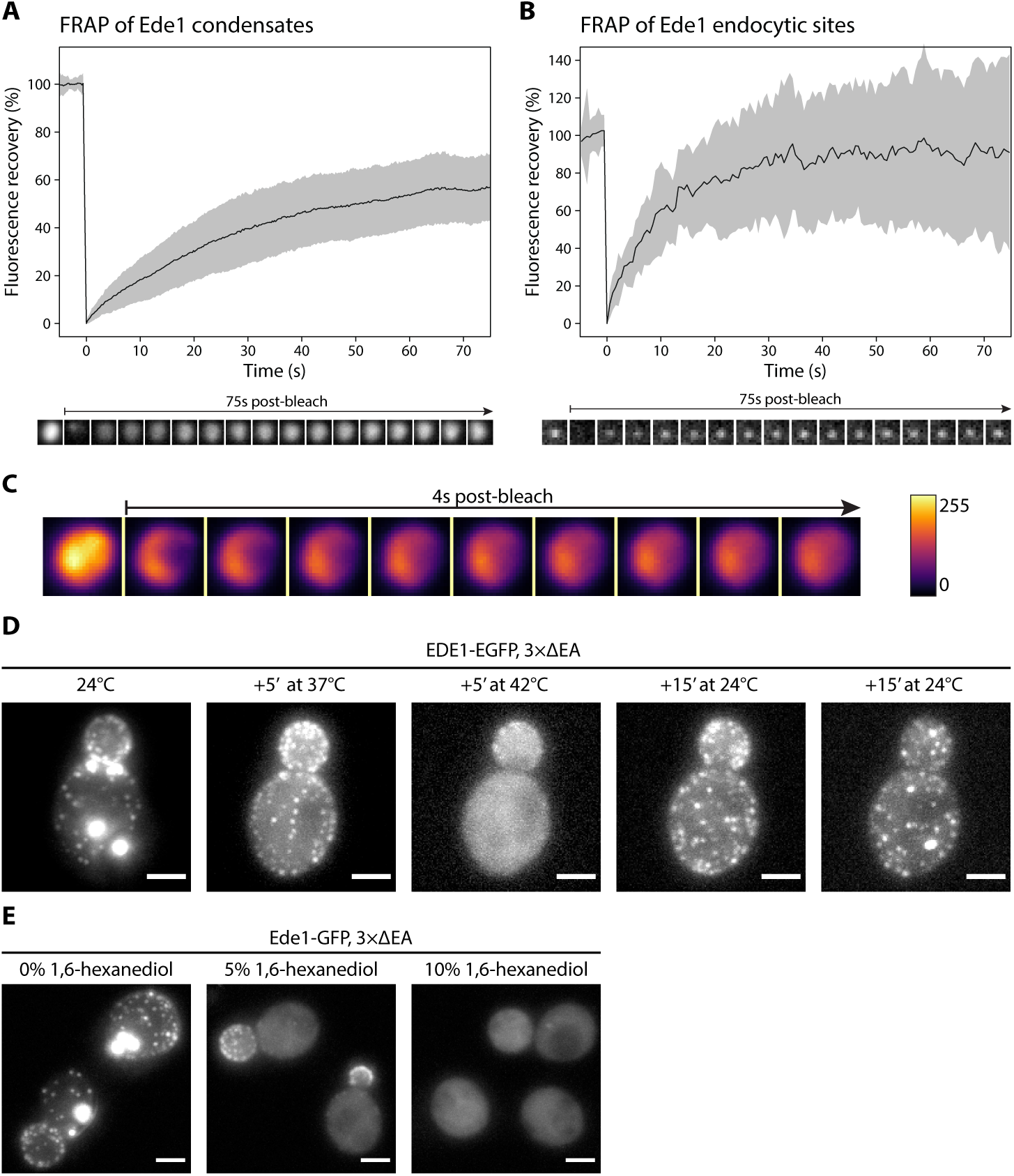
Ede1 condensates and endocytic patches exchange molecules with the cytoplasm and respond to temperature changes. (A, B) Fluorescence recovery after photobleaching of Ede1-EGFP condensates in 3×ΔEA cells (A) and endocytic sites in normal cells (B). Plots show mean fluorescence recovery ± SD. Plot in panel A represents pooled data from four independent experiments (n = 36) and in panel B, from three independent experiments (n = 14). Time series of representative experiments are shown below each plot. In both montages, each frame is 1 μm×1 μm. (C) Time series of a partial bleaching of a condensate in EGFP-Ede1 overexpression strain. A perceptually uniform colour lookup table has been applied to highlight the changes in intensity. Intensity gradient shown for reference. Each frame is 1.5 μm×1.5 μm. (D) Ede1-EGFP was imaged in 3×ΔEA cells at different temperatures. Cells were grown and imaged at 24 °C. Afterwards, the temperature was raised to 37 °C and 42 °C and returned to 24 °C for the indicated amounts of time. Maximum z-stack projections. (E) Representative cells after 5 minute treatment with indicated concentrations of 1,6-hexanediol. Maximum z-stack projections. All scale bars: 2 μm. **Figure 2–video 1.** Multiple partial bleaching experiments in Ede1 overexpression cells. Scale bar: 1 μm.

We then examined the recovery of Ede1 in normal endocytic sites, photobleaching them during TIRF imaging. We found that Ede1-EGFP also turns over fast at endocytic sites (half-time of 7.8 s and mobile fraction of 91 %). The turn over at the endocytic sites was faster and the mobile fraction higher than in the condensates.

We also performed experiments where we only partially bleached the Ede1 condensates to visualize possible internal diffusion of fluorescent molecules. For these experiments we used Ede1-EGFP overexpression cells, in which the condensates are bigger than in the 3×ΔEA cells. When a subregion of an endocytic condensate was bleached, the fluorescence in the bleached and unbleached regions equalized within seconds (Figure 2C, Figure 2–video 1).

The formation of phase-separated condensates depends on protein concentration and can be affected by environmental factors such as temperature (Franzmann et al., 2018; Molliex et al., 2015; Nott et al., 2015). When the 3×ΔEA cells cultured at 24 °C were incubated at 37 °C for 5 minutes, the condensates dissolved while the endocytic sites persisted (Figure 2D). When the temperature was raised to 42 °C, Ede1 signal became entirely diffuse. This effect was reversible for both the endocytic sites, which reformed after a couple of minutes, and the condensates, which reappeared within 30 minutes after return to 24 °C.

1,6-hexanediol is an aliphatic alcohol that disrupts weak protein-protein interactions (Patel et al., 2007) and is used to distinguish between solid and liquid protein aggregates (Kroschwald et al., 2015). Both Ede1 condensates and endocytic patches rapidly disappeared in 3×ΔEA cells upon 1,6-hexanediol treatment (Figure 2E). Interestingly, endocytic patches only fully dissolved at higher 1,6-hexanediol concentrations than Ede1 condensates.

Taken together our results show that Ede1 both in the condensates and at the endocytic sites behaves in a highly dynamic, liquid-like manner.

### Ede1 condensates recruit other endocytic proteins

We then imaged double-tagged strains to test whether other endocytic proteins colocalize with Ede1 condensates in the 3×ΔEA background (Figure 3A, B). We found that the condensates contain multiple early (Syp1, Ent1, Sla2) and late (End3, Pan1, Sla1) coat proteins, as well as the actin nucleation-promoting factor Las17, known to physically interact with End3/Pan1/Sla1 complex (Feliciano and Di Pietro, 2012; Sun et al., 2015).

**Figure 3.**
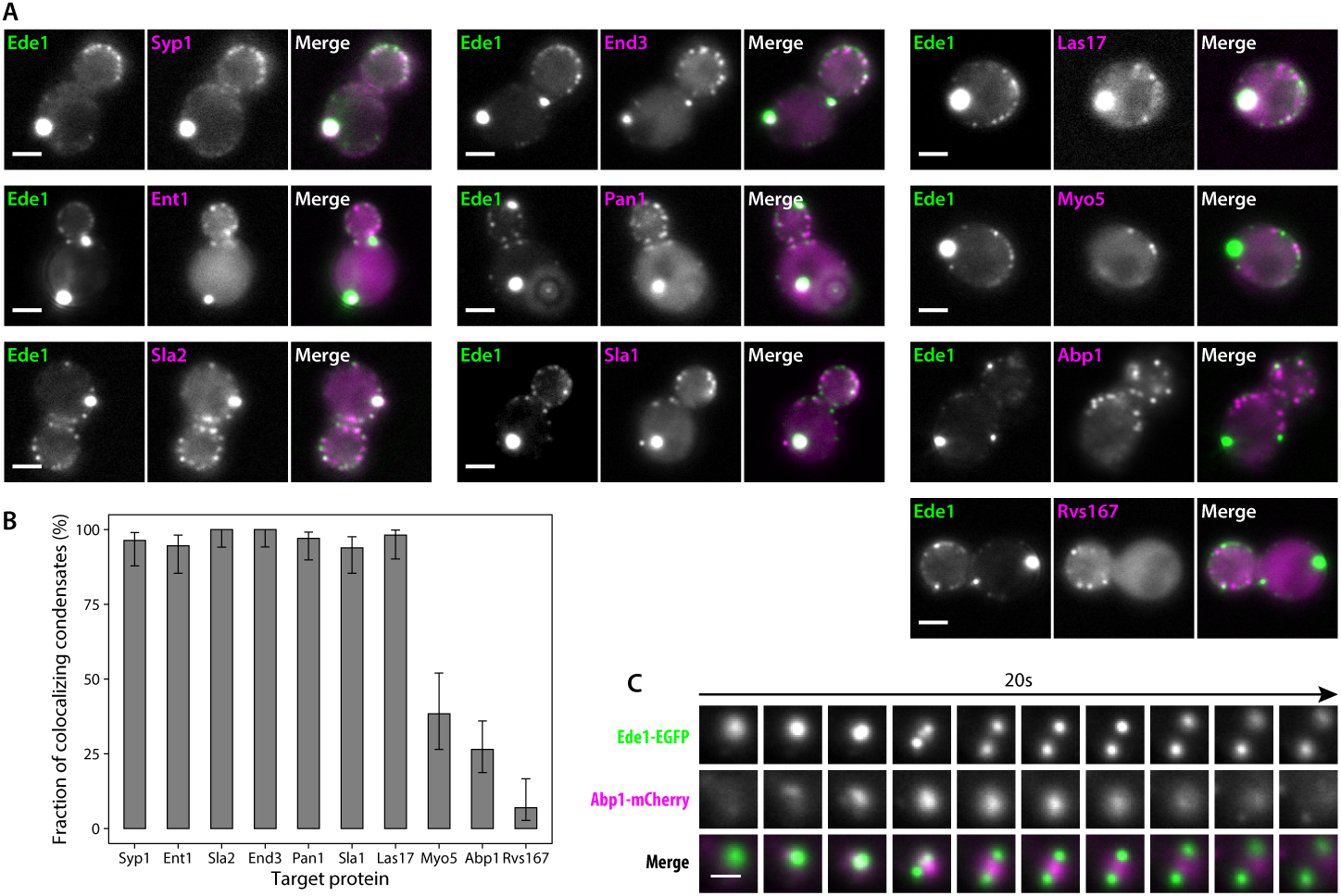
Endocytic condensates recruit many proteins. (A) Images of representative cells expressing Ede1-EGFP and indicated endocytic proteins tagged with mCherry in 3×ΔEA background. (B) Fraction of Ede1-EGFP condensates that colocalized with mCherry puncta in each strain from panel A. Percentages from single experiments with number of analyzed condensates between 52 and 98. Error bars show 95 % binomial confidence intervals estimated using the Wilson score method. (C) Montage from timelapse imaging of Ede1-EGFP and Abp1-mCherry during apparent fission of an Ede1 condensate. **Figure 3–video 1.** A 2-minute movie of Ede1-EGFP (green) and Abp1-mCherry (magenta) in 3×ΔEA background showing repeated transient localization of Abp1 to Ede1 condensates, and an example of Abp1 recruitment coinciding with condensate fission and subsequent fusion. Scale bar 2 μm.

Three proteins Myo5, Abp1 and Rvs167 whose arrival overlaps with actin polymerization at the endocytic sites (Sun et al., 2006) localized to a minority of the condensates (38 %, 27 % and 7 %, respectively). We further examined the interaction of condensates and Abp1 using timelapse imaging and found that Abp1-mCherry patches localized transiently on the condensates (Figure 3C), which explains the partial colocalization. Abp1-mCherry was also present whenever the fission of condensates occurred (Figure 3C, Figure 3–video 1). The fact that fission events coincide with the appearance of Abp1 suggests that the condensates trigger the polymerization of actin filaments which can exert force and cause the fission.

Overall, the colocalization experiments showed that the endocytic condensates are complex, and that the proteins contained within them are at least partially functional as they can recruit their interaction partners and catalyze cycles of assembly and disassembly of actin.

### Ede1 core region is necessary and sufficient for phase separation

Ede1 is a 1381 amino acid long, multi-domain protein (Figure 4A). Its N-terminal region contains three Eps15-homology (EH) domains that interact with asparagine-proline-phenylalanine (NPF) motifs found on endocytic adaptors such as Ent1/2, Yap1801/2 and Sla2 (Maldonado-Báez et al., 2008). Such repeats of domains interacting with linear motifs are known to promote liquid-liquid phase separation (Banjade and Rosen, 2014; Li et al., 2012). The EH domains are followed by a proline-rich region and a coiled-coil domain (Lu and Drubin, 2017; Reider et al., 2009). The C-terminal half of Ede1 contains a Syp1-interacting region (Reider et al., 2009) and a ubiquitin-associated (UBA) domain.

**Figure 4.**
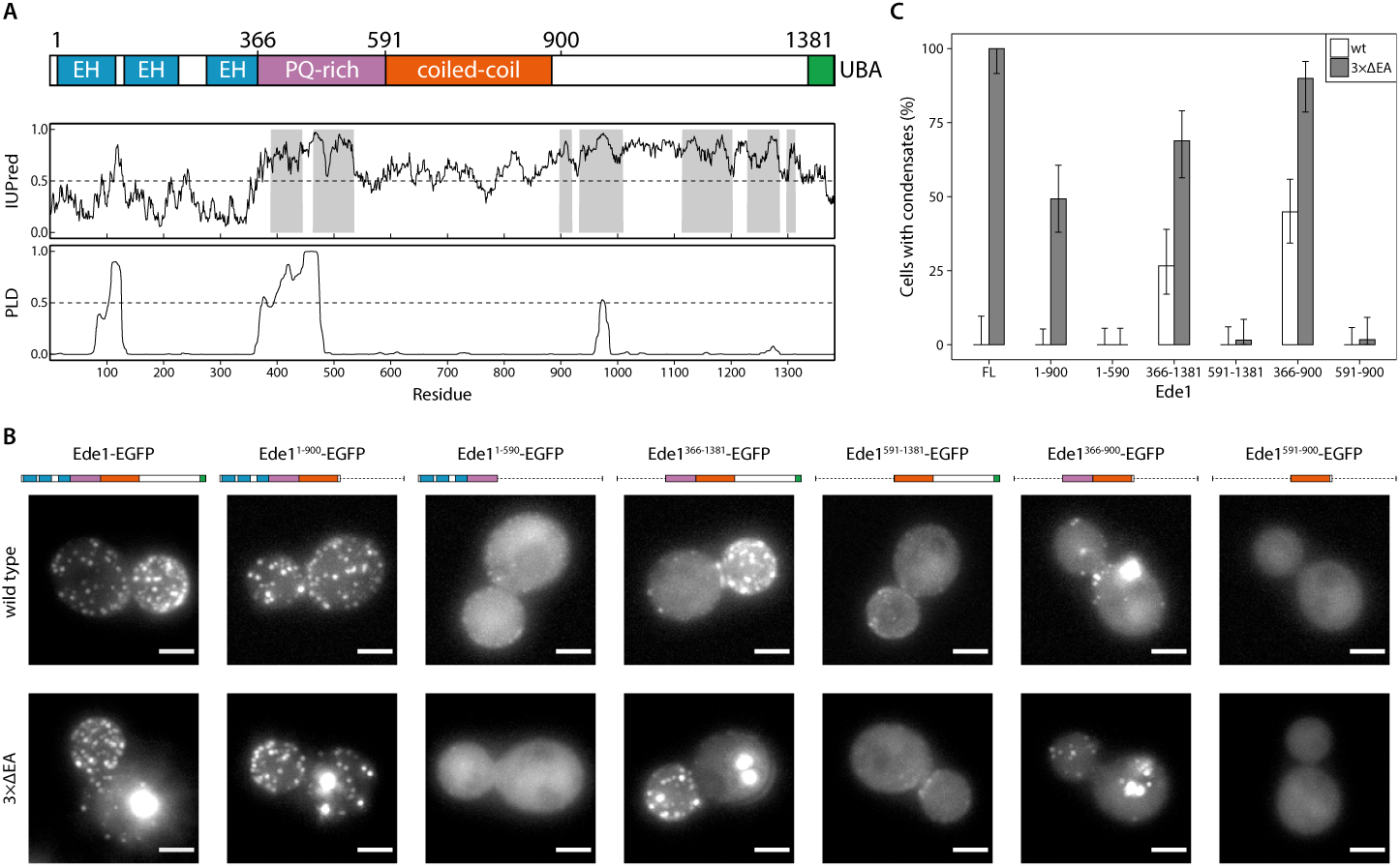
The core region of Ede1 is necessary and sufficient for condensate formation. (A) The domain structure of Ede1 and prediction of disordered and prion-like regions. EH, Eps15-homology domain; UBA, ubiquitin-associated domain. Domains are drawn to scale according to UniProt entry P34216, and numbers above mark domain boundaries used in our constructs. The top plot represents IUPred2a disorder prediction score, with the shaded areas predicted to be disordered by MobiDB-lite consensus method. PLD score indicating the prion-like character was calculated using PLAAC. (B) Representative cells expressing full-length Ede1 and its truncation mutants in wild-type and 3×ΔEA backgrounds. All constructs are C-terminally tagged with EGFP. Maximum intensity projections of 3D volumes, scale bars: 2 μm. (C) The fraction of cells in strains from panel B that show condensates of the imaged construct. Number of cells for each strain between 50 and 78, error bars represent binomial confidence intervals estimated using the Wilson score method.

We noticed that the proline-rich region contains a high number of asparagine and glutamine residues, a hallmark of prion-like domains (Alberti et al., 2009) which are proposed to regulate phase separation (Franzmann and Alberti, 2019; Franzmann et al., 2018). We used PLAAC (Lancaster et al., 2014), a web-based version of the algorithm used by Alberti et al. (2009) to detect prion candidates in yeast proteome, to analyze the Ede1 sequence (Figure 4A). The algorithm detected a 99 amino acid long prion-like sequence between amino acids 374 and 472, suggesting that this region could also be involved in the phase separation of endocytic proteins. We also consulted IUPred2a (Mészáros et al., 2018) and MobiDB-lite (Necci et al., 2017) algorithms to predict intrinsically disordered regions (IDRs) in Ede1. 36 % of Ede1 is predicted to be disordered; the unstructured regions are contained within the proline- and glutamine-rich region, and between the coiled-coil and the UBA domain.

To understand which features of Ede1 mediate phase-separation, we expressed a series of truncations of Ede1 in both wild-type and 3×ΔEA backgrounds, and analyzed their localization to condensates and endocytic sites (Figure 4B,C). In our 3×ΔEA background, the N- and C-terminal regions were dispensable for condensate formation. Surprisingly, Ede1 core region consisting of amino acids 366-900 (the unstructured PQ-rich region and the coiled-coil domain) localized to large condensates in both the 3×ΔEA and wild-type backgrounds. It was also the minimal construct to form these condensates, as constructs containing only the coiled-coil or the PQ-rich region showed diffuse cytoplasmic localization.

For the wild-type background, our results are largely consistent with previously published results about the localization of Ede1 mutants (Boeke et al., 2014; Lu and Drubin, 2017). Namely, the coiled-coil domain was necessary for Ede1 to assemble into endocytic sites, while the N- and C-terminal parts of Ede1 were individually dispensable.

### The functional significance of Ede1 core region

To test the role of the core region of Ede1 in endocytosis, we created EGFP tagged Ede1 mutants with internal deletions of amino acids 366-590 (Ede1^ΔPQ^), 591-900 (Ede1^ΔCC^) and 366-900 (Ede1^ΔPQCC^). Ede1^ΔPQCC^ failed to localize to endocytic sites, whereas the two single-domain Ede1^ΔPQ^ and Ede1^ΔCC^ deletion mutants were still punctate, but more diffuse compared to full-length Ede1-EGFP (Figure 5A).

**Figure 5.**
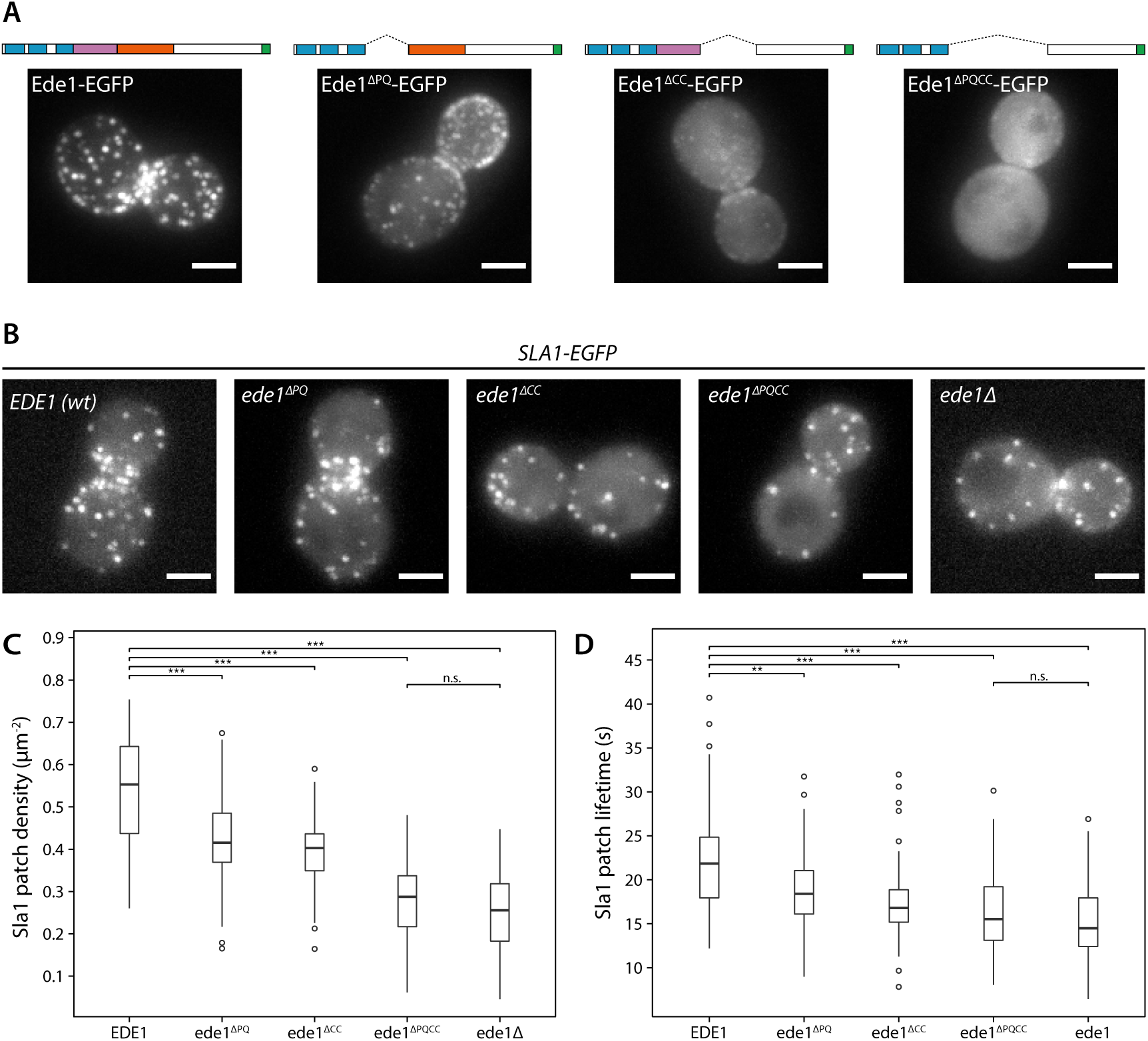
Ede1 features necessary for phase separation are also crucial for its function. (A) Representative cells expressing full-length Ede1 and three internal Ede1 deletion constructs: Ede1^ΔPQ^ (Δ366-590), Ede1^ΔCC^ (Δ591-900) and Ede1^ΔPQCC^ (Δ366-900) tagged with EGFP. (B) Micrographs of representative budding cells expressing Sla1-EGFP and indicated Ede1 internal deletion constructs. (C, D) The number of patches of late coat protein Sla1 per surface area (33-56 yeast cells per strain) and Sla1 lifetimes (49-61 patch trajectories per strain). Scale bars: 2 μm.

Next, we tested if these Ede1 mutants had endocytic defects by using Sla1 as a reporter of the late phase of endocytosis. We tagged Sla1 with EGFP in Ede1 mutant strains (Figure 5B) and quantified the density and lifetimes of endocytic sites (Figure 5C, D). In *ede1Δ* and *ede1*^*ΔPQCC*^ cells, the median number of endocytic events marked by Sla1-EGFP per μm^2^ was reduced by 54 % and 48 %, respectively. Consistent with their effects on Ede1 recruitment, the *ede1*^*ΔPQ*^ and *ede1*^*ΔCC*^ mutations caused intermediate reduction in patch density (25 % and 27 %, respectively). All mutants had significantly less events than the wild type (p < 0.001 in Wilcoxon rank-sum test), and the difference between *ede1Δ* and *ede1*^*ΔPQCC*^ was not statistically significant (p > 0.05).

The Sla1 lifetimes were likewise affected by the deletion of Ede1 core regions. In *ede1Δ*, Sla1-EGFP lifetime was decreased by 34 % and in *ede1*^*ΔPQCC*^, by 29 %. The deletions of individual regions again showed intermediate defects (16 % and 23 % for Ede1^ΔPQ^ and Ede1^ΔCC^, respectively). All mutants were significantly different than the wild type (p < 0.001, except for *ede1*^*ΔPQ*^ where p < 0.01). The full Ede1 deletion was not significantly different from *ede1*^*ΔPQCC*^ (p > 0.05).

Taken together, our results show that the core region is essential for Ede1 to promote efficient endocytosis and to regulate the timing of coat maturation.

### Ede1 core region facilitates localization of other early proteins

In *ede1Δ* cells many of the early endocytic proteins fail to localize to endocytic sites (Carroll et al., 2012; Stimpson et al., 2009). We therefore visualized the localization of early proteins in Ede1 mutants lacking the core region.

Different proteins were affected by the Ede1 core deletions in different ways, consistent with the work done on *ede1Δ* mutants (Figure 6). The localisation of Apl1 (β-subunit of the AP-2 complex) was the most severely disrupted. Apl1-EGFP patches were less defined in Ede1^ΔPQ^ background, and undetectable in Ede1^ΔCC^ or Ede1^ΔPQCC^ cells. In these cells, Apl1-EGFP signal was dispersed in the cytoplasm, with a faint presence around the bud neck. Syp1 and Yap1801 remained localized to the membrane in all of the mutants, but the signal became more diffuse along the membrane and the patches less defined. This effect was the strongest for Ede1^ΔPQCC^, with intermediate effects in Ede1^ΔPQ^ and Ede1^ΔCC^ cells. However, Ent1 and Sla2 were were still assembled into endocytic patches in all the Ede1 mutants. Taken together, our results indicate that the Ede1 core region is essential to concentrate early endocytic proteins.

**Figure 6.**
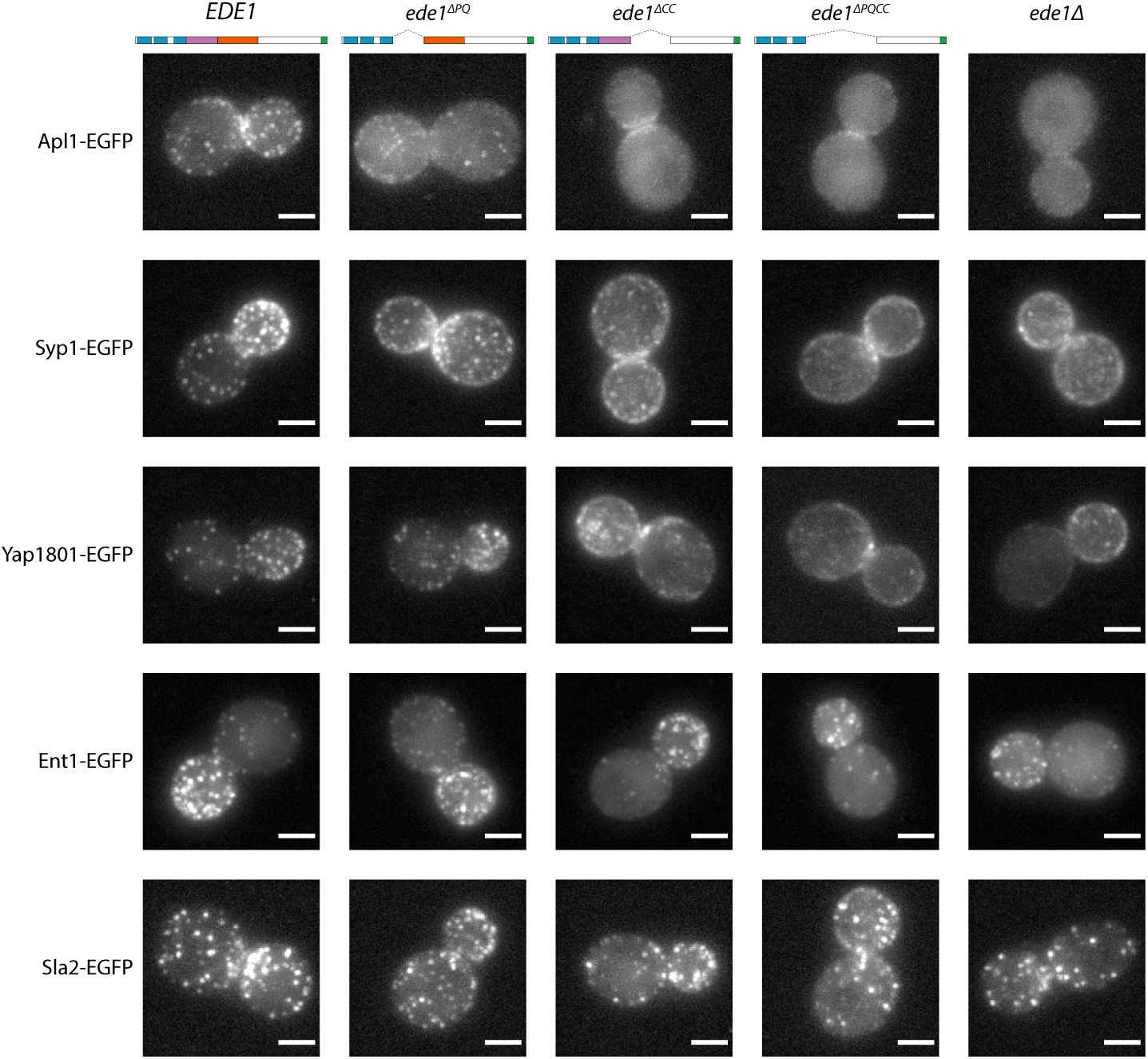
Ede1 core deletion mutants are defective in early protein localization. Maximum-intensity projections of 3D volumes are shown for representative cells with different early proteins tagged with EGFP. The strains express Ede1 species indicated at the top. Scale bars: 2 μm.

### Ede1 core region can cluster a heterologous lipid-binding protein

We hypothesized that the phase separation mediated by Ede1 core region is able to cluster membrane associated proteins. To test our hypothesis, we fused the Ede1 core to a diffusely membrane-bound protein. We created a GFP-Ede1^366-900^-2×PH(PLCγ) construct based on a PI(4,5)P_2_ probe developed by Stefan et al. (2002). The original GFP-2×PH(PLCγ) construct is distributed homogeneously on the plasma membrane, while GFP-Ede1^366-900^ alone localized to bright intracellular condensates. In contrast, the fusion construct localized to the plasma membrane, forming puncta that resembled endocytic sites (Figure 7A).

**Figure 7.**
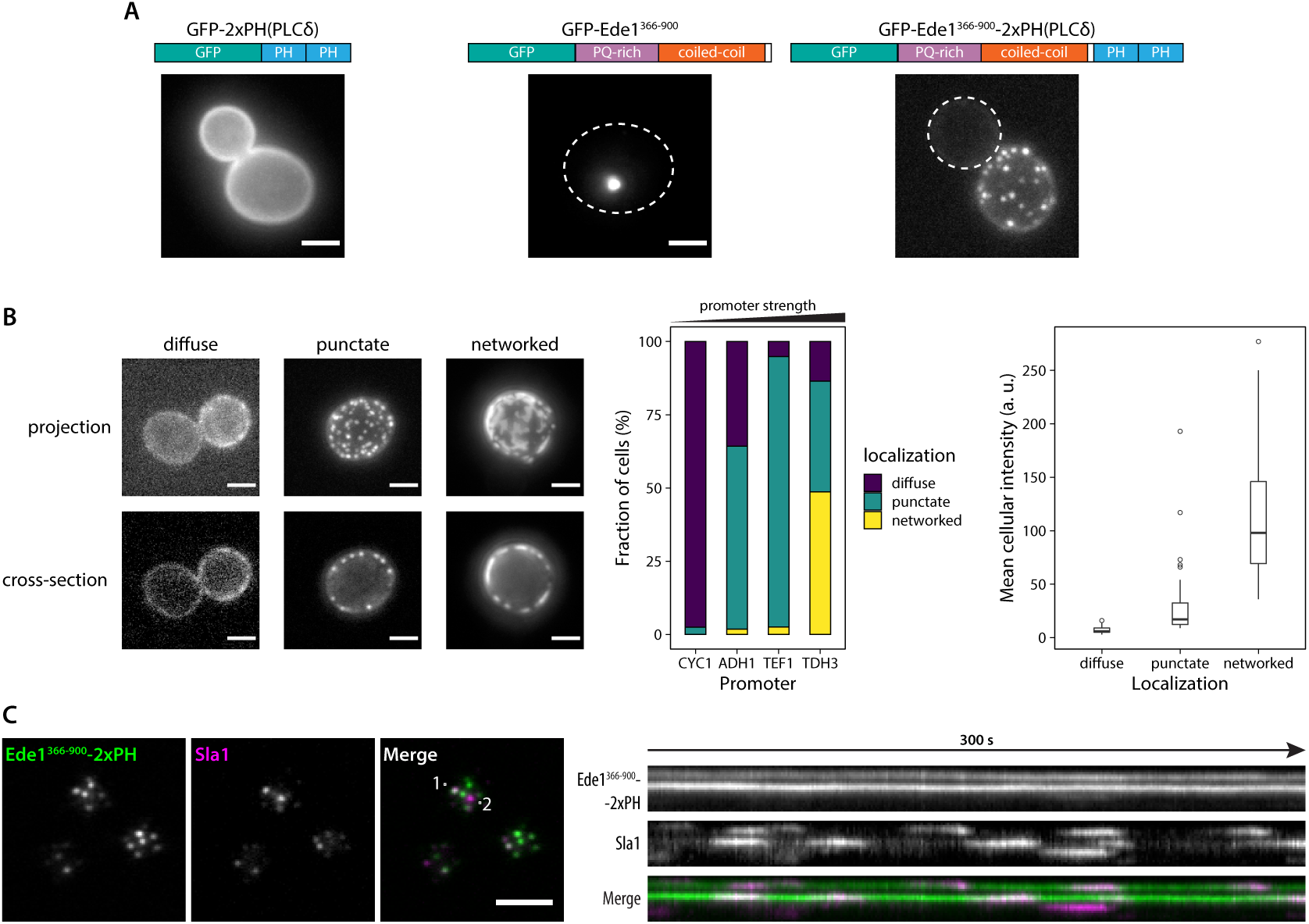
Fusion with Ede1 core region changes the distribution of a PI(4,5)P_2_ probe. (A) Maximum projections of cells expressing GFP-2×PH, GFP-Ede1^366-900^, and GFP-Ede1^366-900^-2×PH. All constructs expressed from a yeast centromeric plasmid under the control of TDH3 promoter. White dotted line shows cell outline. (B) Signal from cells expressing GFP-Ede1^366-900^-2×PH was classified as ‘diffuse’, ‘punctate’ or ‘networked’. Left: class examples; middle: percentage of cells falling into each class when the construct is expressed under the control of different promoters; right: mean cellular fluorescence intensity for each class, after background subtraction, from micrographs obtained under the same imaging conditions. (C) Time series of cells expressing GFP-Ede1^366-900^-2×PH and Sla1-mCherry were acquired using TIRF microscopy. Left: single frame from a representative movie. Points labeled ‘1’ and ‘2’ mark the top and bottom of the kymograph (right). Scale bars: 2 μm. All cells in this figure: *SLA1-mCherry::KANMX4, ede1Δ::natNT2*. **Figure 7–video 1.** The 5-minute TIRF movie of GFP-Ede1^366-900^-2×PH (green) and Sla1-mCherry (magenta) represented in panel C. Scale bar: 2 μm **Figure 7–video 2.** A 3-minute equatorial plane movie of GFP-Ede1^366-900^-2×PH (green) and Sla1mCherry (magenta) showing inward movement of Sla1-mCherry patches. Scale bar: 2 μm.

We also noticed subpopulations of cells with different localization patterns of the construct (Figure 7B). We speculated that the variable patterns were caused by heterogeneity in protein expression level due to plasmid copy number variation. To test that hypothesis, we expressed GFP-Ede1^366-900^-2×PH(PLCγ) from centromeric plasmids containing four different promoters of increasing strength (Mumberg et al., 1995). We classified the localization of the construct in these cells as ‘diffuse’, ‘punctate’ or ‘networked’. We found that the tendency to cluster into different patterns correlated with promoter strength and the expression level. Low expressing cells had more diffuse localization of the construct, and separated into puncta or well-separated regions as the concentration increased.

Intriguingly, the puncta formed by the GFP-Ede1^366-900^-2×PH(PLCγ) construct were stable over long imaging periods, but dynamically recruited the late coat protein Sla1 (Figure 7C, Figure 7–video 1). Sla1-mCherry persisted at these sites with similar lifetimes as during normal endocytosis and showed inward movement (Figure 7–video 2), indicating functional vesicle budding.

These results show that directing the phase-separating Ede1 core domain to the plasma membrane is sufficient to create puncta on the membrane in a concentration-dependent manner. Surprisingly, these long lived artificial sites can repeatedly initiate endocytic events and thereby function as artificial endocytic hot-spots.

## Discussion

Our results indicate that the core region of Ede1 can phase separate in cells, and that the same region is also necessary for Ede1’s function in promoting the initiation of endocytic events. We also demonstrated that the core region of Ede1 fused to a lipid-binding domain can condense on the plasma membrane in a concentration dependent manner. These findings link endocytic assembly to the phenomenon of protein phase separation.

### Are the Ede1 condensates phase separated liquids?

Ede1 forms liquid-like condensates under conditions where the stoichiometry between the Ede1 and endocytic adaptor proteins is altered, such as overexpression of Ede1, or deletion of three Ede1 interacting adaptors (Boeke et al. 2014, Figure 1). It is often challenging to demonstrate that a protein forms liquid condensates *in vivo*, but there are several commonly applied criteria (reviewed by Alberti et al. 2019). These criteria include direct visualization of liquid-like behaviors, such as fusion, shearing under flow and relaxation into spherical droplets. A fast turnover of molecules, which can be demonstrated with FRAP experiments, is another hallmark of liquid-like condensates. Furthermore, the dependency of the condensate assembly on physical parameters such as protein concentration and temperature are frequently used criteria. Susceptibility to disruption by 1,6-hexanediol has often been used to differentiate solid and liquid-like condensates (Kroschwald et al., 2015).

We showed that the Ede1 condensates undergo apparent fusion and fission events, the latter caused likely by force generated by polymerizing actin filaments (Figure 3C, Figure 3–video 1). The Ede1 molecules exchange rapidly between the condensate and the cytosolic pool (Figure 2A) and, importantly, Ede1 molecules also diffuse fast within the condensates (Figure 2C). The formation of the Ede1 condensates is dependent on the cellular concentration of Ede1 (Figure 1D). The condensates dissolve rapidly and reversibly in response to temperature changes (Figure 2D, and are sensitive to treatment with 1,6-hexanediol (Figure 2E).

In agreement with our findings in yeast, Day et al. (2019) have recently shown that Eps15, the mammalian homologue of Ede1, can phase separate *in vitro*. Taken together, these data suggest that Ede1 has the ability to form phase separated condensates in cells under conditions that are close to normal physiological conditions.

### Are the early endocytic sites phase separated condensates?

Under normal conditions Ede1 does not form large condensates, but only transiently assembles at endocytic sites. We hypothesize that Ede1 molecules form a liquid phase at the nascent endocytic sites, concentrating other early endocytic proteins and driving the initiation of an endocytic event. Three of the criteria we used to characterize the liquid-like nature of the Ede1 condensates could be applied to Ede1 at the normal endocytic sites. Namely, we showed that the Ede1 molecules turn over fast between the endocytic site and the cytosolic pool (Figure 2A,B), that the assembly of Ede1 at the endocytic sites is sensitive to temperature (Figure 2D), and that it can be disrupted by 1,6-hexanediol (Figure 2E).

Interestingly, both the temperature and the 1,6-hexanediol concentration that blocked the assembly of Ede1 at endocytic sites were higher than those that disassembled the Ede1 condensates. The dimensionality of a system plays an important role in determining its phase behaviour, and it has been demonstrated *in vitro* that in a two-dimensional system the threshold concentration for phase separation can be an order of magnitude lower than in a three-dimensional system (Banjade and Rosen, 2014). Our results show that the two-dimensional, membrane-limited endocytic sites are the preferred structures over the cytosolic condensates. This suggests that the cells might operate in a concentration regime sufficient to promote two-, but not three-dimensional phase separation of endocytic proteins.

The size of the endocytic sites below the resolution limit of light microscope makes it challenging to detect the possible liquid-like behaviors, such as fusion. However, superresolution imaging of Ede1 and other endocytic proteins in fixed cells revealed that Ede1 forms larger and more amorphous structures than clathrin and clathrin adaptor proteins (Mund et al., 2018).

Furthermore, the observation that the Ede1 condensates efficiently recruit early endocytic proteins points to a functional similarity between the condensates and the normal endocytic site. Finally, the fact that the same core region of Ede1 is essential both for the formation of the condensates and for the assembly of Ede1 at the endocytic sites suggests that the assembly mechanisms of the condensates and the endocytic sites are related. We therefore propose that Ede1 promotes endocytic assembly through its phase separation activity.

### Molecular mechanisms of Ede1’s function

The N- and C-terminal regions of Ede1 mediate the known protein-protein interactions between Ede1 and endocytic adaptor proteins, while the core region mediates Ede1’s self-assembly (Aguilar et al., 2003; Boeke et al., 2014; Reider et al., 2009). Both the terminal protein-protein interactions and the self-interacting core are needed for the function of Ede1 in endocytic initiation (Lu and Drubin, 2017). We showed here that the core region of Ede1 is sufficient and necessary for Ede1’s phase separation in cells, and that it is also essential for the function of Ede1 in initiating endocytic events. The adaptor interactions recruit Ede1 to the plasma membrane, increasing its local concentration and likely promoting its phase separation.

In addition, Ede1 can interact with transmembrane endocytic cargo proteins, either directly via its ubiquitin binding domain or indirectly via adaptor proteins. In this way the cargoes could also increase the local concentration of Ede1 and thereby promote its phase separation. This could be the molecular mechanisms by which cargo molecules promote endocytosis.

### Building artificial endocytic sites

To test the idea that phase separation could promote the initiation of endocytosis we created a chimeric protein that combined the phase separating core regions of Ede1 with membrane associating PH domains (Figure 7). The PH domains on their own bind diffusely on the plasma membrane, whereas the Ede1 core region mainly localizes to cytosolic condensates. We hypothesized that the PH domains would bring the chimeric protein to the plasma membrane and that the Ede1 core region would phase separate to generate high local concentrations of the protein. Indeed, the chimeric construct localized to the plasma membrane in a punctate pattern that resembles the distribution of endocytic sites. The artificial sites were, however, long-lived compared to the normal endocytic sites, which disassemble after vesicle budding. The artificial sites are likely uncoupled from the mechanisms that normally trigger the disassembly of the endocytic proteins. The artificial sites were also not enriched in the growing bud of the yeast cell like the normal endocytic sites are. Their initiation is likely uncoupled from the factors, such as cargoes, that normally determine the distribution of endocytic events. In addition, because they are long-lived, the artificial sites may sequester the chimeric protein and thus prevent the formation of new sites in the bud.

The distribution pattern of the chimeric protein on the plasma membrane depended on its cellular concentration. At low levels the protein remained diffuse on the plasma membrane, while at intermediate levels it formed a punctate pattern typical of endocytic proteins. At higher expression levels, however, we observed a networked pattern on the membrane. Such patterns are reminiscent of those observed during decomposition in the spinodal concentration range (Alberti et al., 2019; Berry et al., 2018).

Surprisingly, the artificial sites formed by the chimeric protein could initiate vesicle budding events. As the artificial sites are not disassembled with vesicle budding they could initiate repeated budding events thus generating endocytic ‘hot spots’, which are not normally seen in yeast cells. Since the core region of Ede1 does not have known interactions with other endocytic proteins, the mechanisms by which the chimeric molecule can recruit other endocytic proteins to trigger vesicle budding is not clear. An interesting possibility is that other endocytic proteins phase separate together with Ede1.

### Other roles for phase separation in endocytosis

Intrinsically disordered and prion-like domains are an established determinant of the phase separation of proteins (Franzmann et al., 2018; Patel et al., 2015), as are multivalent interactions of linear motifs and folded recognition domains (Banani et al., 2016; Li et al., 2012). In the case of Ede1, the coiled-coil and disordered domains are important for its phase separation properties. Many other endocytic proteins have unstructured (Dafforn and Smith, 2004; Kalthoff et al., 2002) and prion-like (Alberti et al., 2009) regions that may have a tendency to phase separate, and recent studies suggested that the unstructured protein regions can generate force for membrane bending during vesicle budding (Bergeron-Sandoval et al., 2018; Busch et al., 2015; Snead et al., 2017).

Endocytic proteins also form a vast network of structured interactions, such as those between NPF motifs and the EH domains of Ede1, End3 and Pan1, and between multiple proteins containing SH3 domains and proline-rich motifs (Tonikian et al., 2009). In fact, phase separation mediated by SH3–PRM interactions has already been identified as a factor in promoting actin polymerisation in membrane-bound signalling clusters (Banjade and Rosen, 2014; Case et al., 2019b; Su et al., 2016).

Protein phase separation processes may thus play important roles at several stages of clathrin-mediated endocytosis, from the initiation of an endocytic event to the final vesicle budding.

## Materials and methods

### Yeast strains and plasmids

The list of yeast strains and plasmids used in this study is provided in Table S1 and Table S2, respectively. Cells were maintained on rich medium at 24 °C or 30 °C.

C-terminally tagged or truncated mutants were generated via homologous recombination with PCR cassettes as described by Janke et al. (2004). N-terminal truncation and internal domain deletion mutants of Ede1 were generated by first constructing the desired mutant gene in a pET-based plasmid. The mutated Ede1 sequence was then amplified by PCR using primers containing 50bp overlap with 5’ (forward primer) 3’ (reverse primer) UTR sequences of EDE1. The PCR product was transformed into *ede1Δ::klURA3* cells. The transformants were selected for on plates containing 5-fluorootic acid, and confirmed by colony PCR and genomic sequencing.

Plasmids used in Figure 7 were based on pRS426-GFP-2×PH(PLCγ) (Stefan et al., 2002), a kind gift from the Emr lab. The Ede1^366-900^ coding sequence was cloned into this plasmid after the last GFP codon using the SLIC method (Li et al., 2012). GFP-2×PH(PLCγ) and GFP-Ede1^366-900^-2×PH(PLCγ) were then subcloned to a pRS416-based plasmid p416-GPD under the control of TDH3 promoter (Mumberg et al., 1995) using BamHI and SalI restriction sites. GFP-Ede1^366-900^-2×PH(PLCγ) was also subcloned into p416-CYC1, p416-ADH1 and p416-TEF1 (Mumberg et al., 1995) using the same restriction sites.

### Live cell imaging

Yeast cells were grown to OD_600_ between 0.3 - 0.8 at 24 °C in synthetic medium lacking tryptophan, or tryptophan and uracil if required for plasmid maintenance. Cells were attached to cover slips coated with 1 mg ml^−1^ concanavalin A. Still images and 3D stacks were obtained on an Olympus IX81 wide-field microscope equipped with a 100x/NA1.45 objective and an ORCA-ER CCD camera (Hamamatsu), using an X-CITE 120 PC (EXFO) metal halide lamp as the illumination source. The excitation and emission light when imaging EGFP- and mCherry-tagged proteins were filtered through the U-MGFPHQ and U-MRFPHQ filter sets (Olympus).

#### Total internal reflection fluorescence microscopy

All TIRF movies were recorded on an Olympus IX83 wide-field microscope equipped with a 150x/NA1.45 objective and an ImageEM X2 EM-CCD camera (Hamamatsu). 488 nm and 561 nm laser lines were used for illumination of GFP- and mCherry-tagged proteins. Excitation and emission were filtered using a TRF89902 405/488/561/647 nm quad-band filter set (Chroma). Laser angles were controlled by iLas2 (Roper Scientific).

#### Fluorescence recovery after photobleaching

Bleaching of Ede1-EGFP in endocytic condensates (Figure 2A,B) was performed using a custom-built set-up that focuses a 488 nm laser beam at the sample plane. The diameter of the bleach spot was approximately 0.5 μm.

Bleaching of unperturbed endocytic sites (Figure 2C) was performed with a 405 nm laser line controlled by the iLas2 targeting system during simultaneous excitation with 488 nm and 561 nm lasers using the TIRF setup described above. The emission light was collected through a Gemini beam splitter (Hamamatsu) equipped with a Di03-R488/561-t1 dichroic, and FF03-525/50-25 and FF01-630/92-25 emission filters (Semrock).

### Image analysis

General image analysis was performed using the Fiji distribution of ImageJ (Rueden et al., 2017; Schindelin et al., 2012). All display images were corrected for background fluorescence using the rolling ball algorithm of ImageJ, and movies were corrected for photo-bleaching using a custom ImageJ macro.

#### FRAP experiments

FRAP experiments performed on Ede1-EGFP condensates were analyzed according to Phair et al. (2004). Mean fluorescence values were measured from regions of interest representing the background, the cell and the condensate. A custom-written R script was used to subtract background fluorescence, correct for photobleaching and normalize the values between 0 (corrected fluorescence immediately after photobleaching) and 1 (mean corrected fluorescence of 5 s before photobleaching). The recovery curves of individual experiments were aligned to bleach time and averaged. Condensates that showed lateral or axial movement during the acquisition were manually excluded from the averaging. The average was fitted to a single exponential equation from which the mobile fraction and recovery half-time were calculated.

For FRAP experiments performed on native endocytic sites, the background fluorescence was first subtracted from the TIRF images using the ImageJ rolling ball algorithm. EGFP and mCherry fluorescence of single endocytic patches was measured within a circle with a radius of three pixels around the patch centroid position. A custom-written R script was used to calculate the fluorescence recovery much in the same way as for the FRAP of condensates, but no further corrections were made for background signal or imaging-induced photobleaching. To calculate average recovery, we manually selected only events in which Abp1-mCherry signal peaked at least 60 s after bleach time to exclude the effect of Ede1 disassembly at the end of endocytic events.

#### Patch numbers and lifetimes

For estimating the number of patches per membrane area, we analyzed single non-budding cells. The patches were thresholded and counted using a custom Python script. We estimated the cell surface area by measuring the area of the cross-section from maximum intensity projection and calculating surface area of a sphere. For estimating patch lifetimes, we tracked endocytic events using ParticleTracker from the MOSAIC suite (Sbalzarini and Koumoutsakos, 2005) and multiplied trajectory length by the frame rate.

#### Cell classification

To calculate the percentages of Ede1-EGFP condensates colocalizing with mCherry puncta in Figure 3, cells containing Ede1-EGFP condensates in Figure 4 and cells showing different GFP-EDE1^366-900^-2×PH localization patterns in Figure 7, single cells were cropped from imaging fields based on a neutral signal (GFP in the case of Figure 3 and brightfield image for Figure 4 and Figure 7). Next, an ImageJ macro was used to display random images from the dataset and the experimenter would assess the presence of the tested phenotype with no knowledge of which strain was being analysed.

## Acknowledgements

This work was supported by the Swiss National Science Foundation (grants 31003A_163267 and 310030B_182825) and by the NCCR Chemical Biology funded by the SNSF.

We are thankful to all the members of the Kaksonen laboratory, especially Markus Mund, Andrea Picco, and Daniel Hummel for their critical reading of the manuscript. We also thank Jeanne Stachowiak and Kasey Day for their comments and Camilla Godlee for contributions to the early phase of the project.

## Author contributions

M. Kozak and M. Kaksonen designed the experiments. M. Kozak performed the experiments and analysed the data. M. Kozak and M. Kaksonen wrote the manuscript. M. Kaksonen secured the funding.

---

**Table S1.**
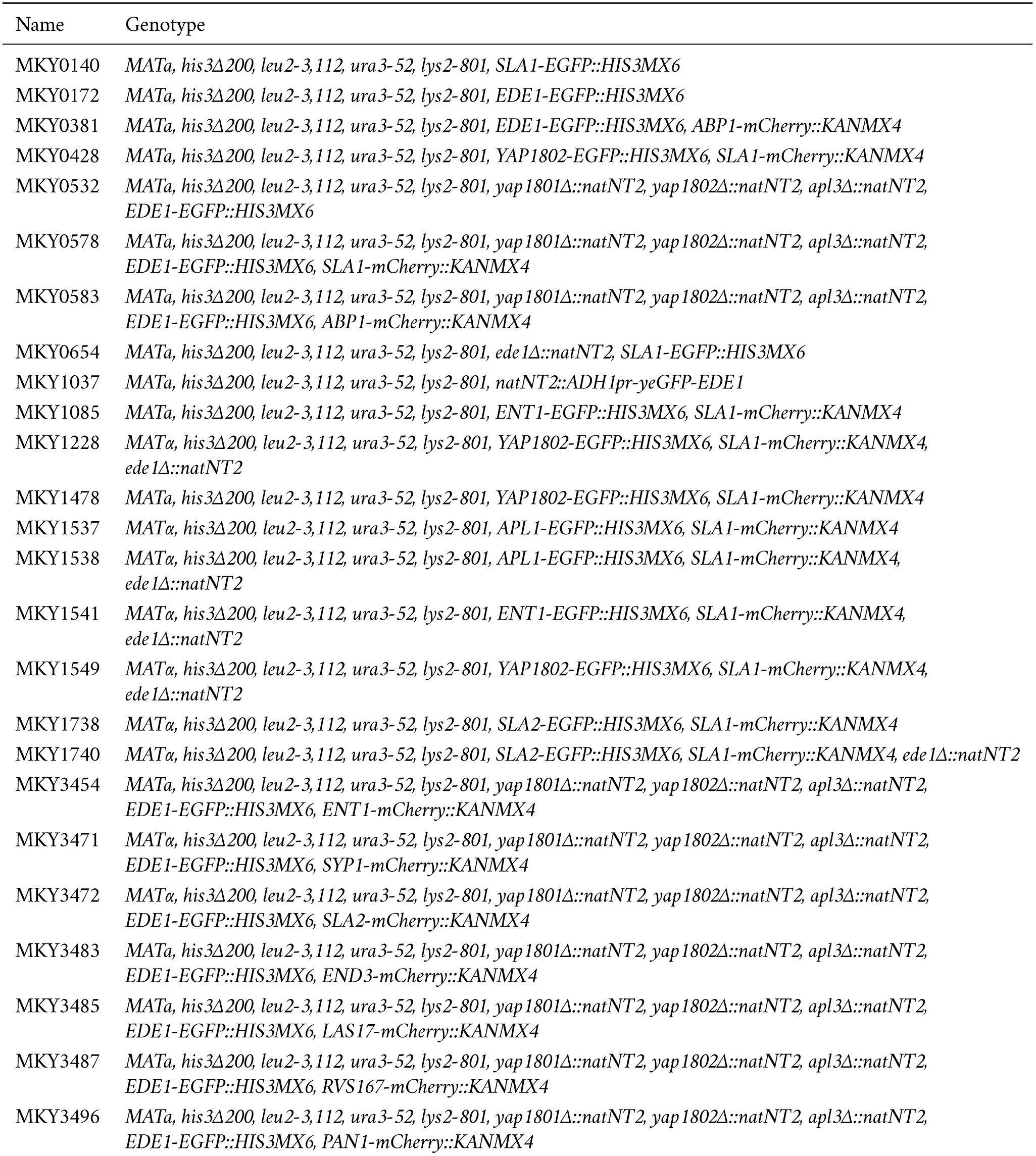
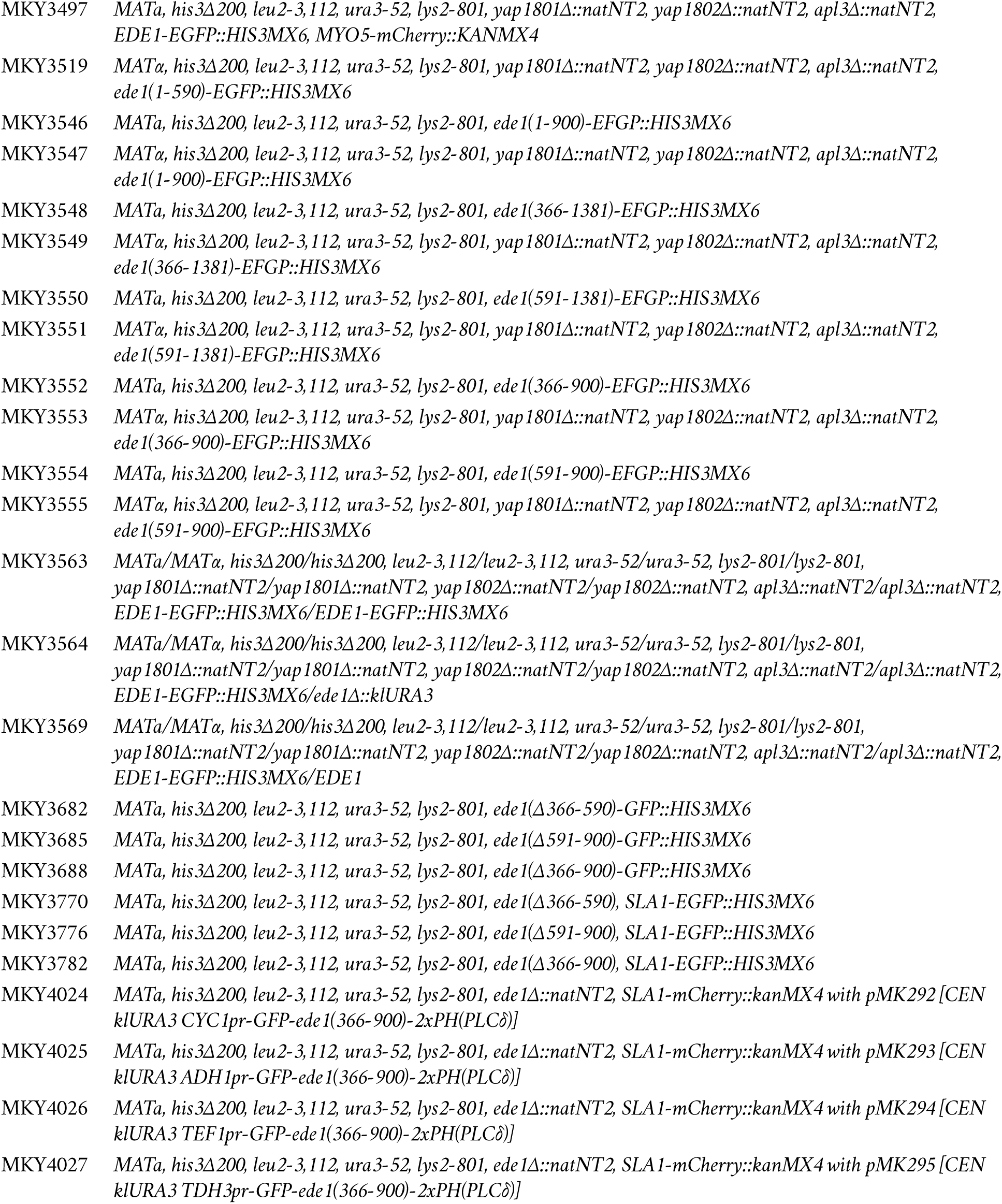
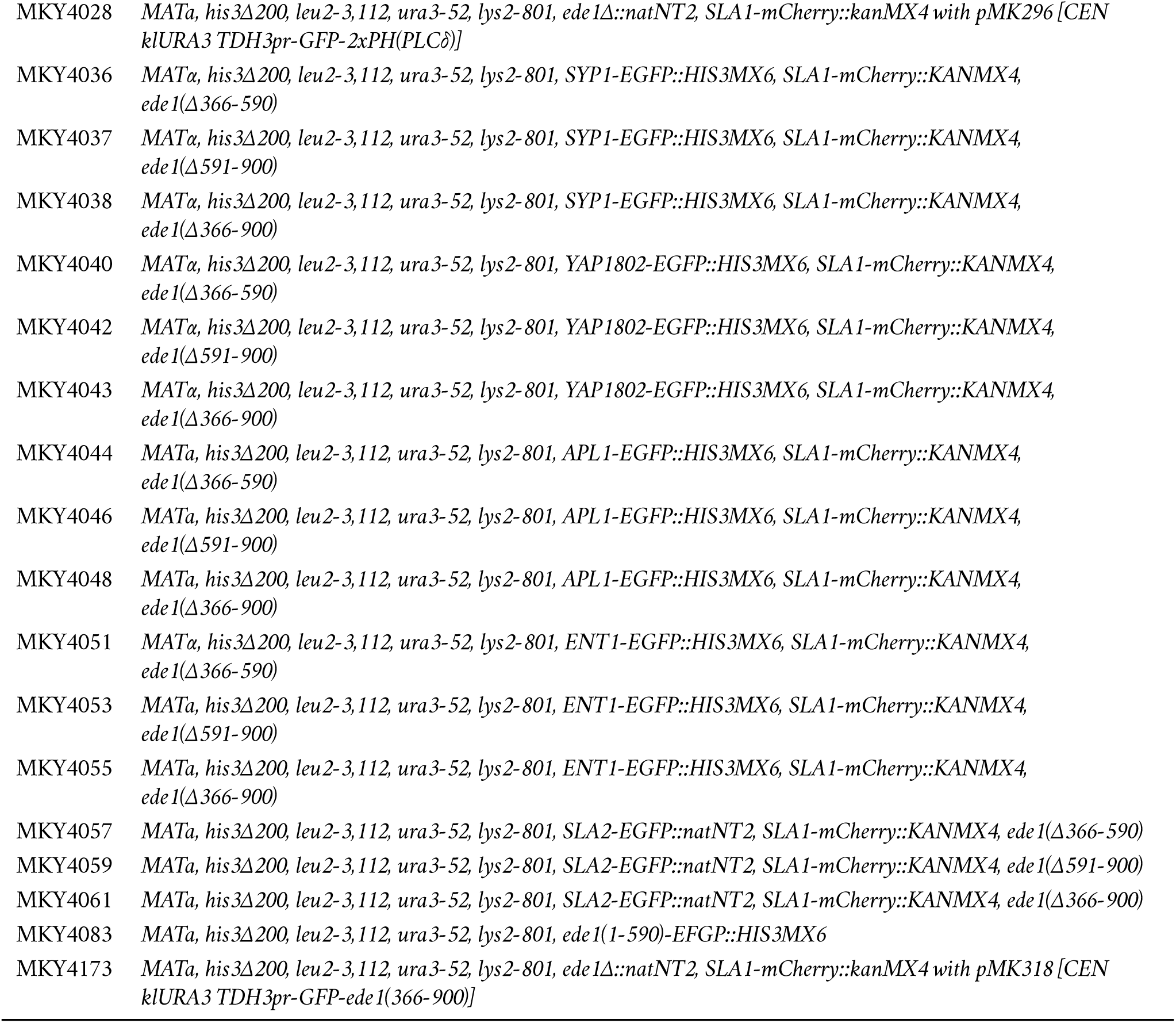
Yeast strains used in this study.

**Table S2.**
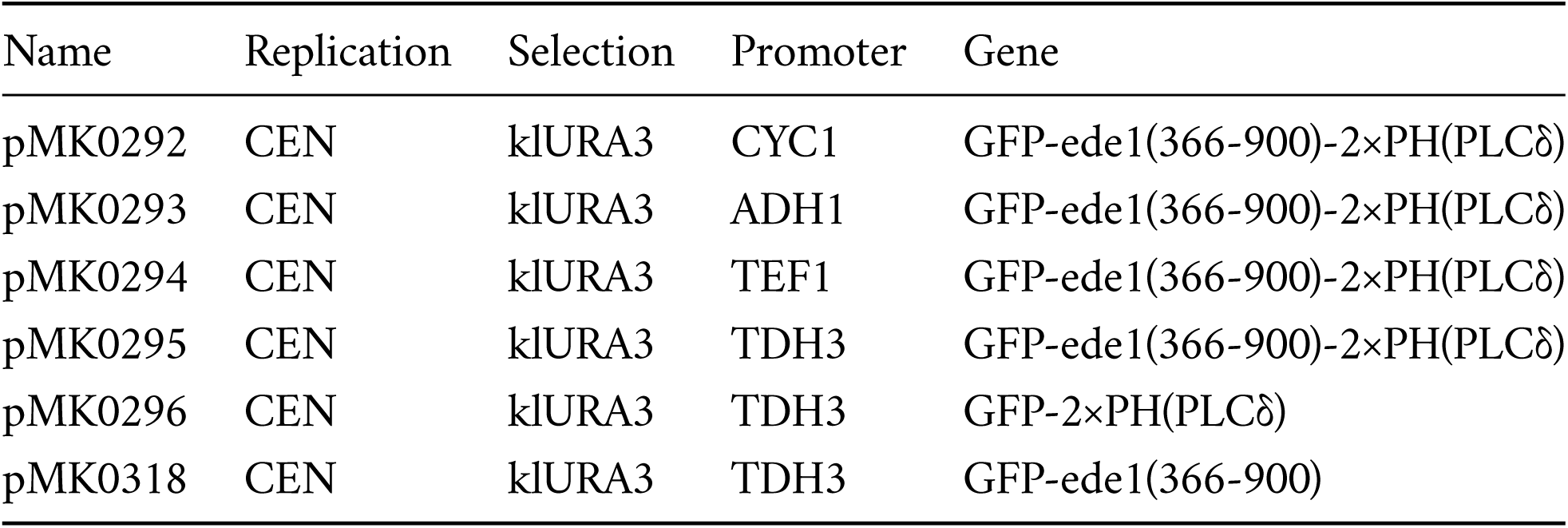
Yeast replicating plasmids used in this study.

